# Long-term history dependence of growth rates of *E. coli* after nutrient shifts

**DOI:** 10.1101/2023.08.22.554350

**Authors:** Dimitris Christodoulou, Avik Mukherjee, Rebekka Wegmann, Adriano Pagano, Varun Sharma, Stephanie Maria Linker, Yu-Fang Chang, Julius Sebastian Palme, Uwe Sauer, Markus Basan

## Abstract

According to a widely accepted paradigm of microbiology, steady-state growth rates are determined solely by current growth conditions^1–3^ and adaptations between growth states are rapid, as recently recapitulated by simple resource allocation models^4^. However, even in microbes overlapping regulatory networks can yield multi-stability or long-term cellular memory. Species like *Listeria monocytogenes*^5^ and *Bacillus subtilis* “distinguish” distinct histories for the commitment to sporulation^6^, but it is unclear if these states can persist over many generations. Remarkably, studying carbon co-utilization of *Escherichia coli*, we found that growth rates on combinations of carbon sources can depend critically on the previous growth condition. Growing in identical conditions, we observed differences in growth rates of up to 25% and we did not observe convergence of growth rates over 15 generations. We observed this phenomenon occurs across combinations of different phosphotransferase (PTS) substrates with various gluconeogenic carbon sources and found it to depend on the transcription factor Mlc.

---

Bacteria occupy diverse ecological niches where they encounter a wide range of growth conditions. Fitness therefore depends on the ability to achieve high growth rates under specific conditions, as well as rapid adaptation to new conditions. To ensure their survival, bacteria need to optimize multiple, sometimes conflicting objectives^7,8^, for example the maximization of growth and the efficient adaptation to new conditions. To better understand, how bacteria achieve this, we followed *E. coli* growth dynamics during shifts from single carbon sources to mixtures of two carbon sources. Initially, we focused on the combination of glucose and acetate because glucose is the preferred carbon source of *E. coli*, supporting high growth rates with glycolytic metabolism^9–11^ and acetate is the primary fermentation product of *E. coli*. Acetate is therefore ubiquitously present in fast growth conditions. As a gluconeogenic carbon source, acetate supports only relatively slow growth and the transition from glucose to acetate leads to long lag phases ^11,12^. For medium shift experiments (Fig. 1A), we first allowed our strain to reach steady-state exponential growth, in two cultures of minimal medium, containing either glucose or acetate as the sole carbon source. After reaching a similar optical density (OD_600_) of about 0.2, we added acetate or glucose such that both tubes contained both glucose and acetate at the same concentration. The concentrations of glucose and acetate used, far exceed total consumed glucose or produced acetate^12^. Cultures were grown until an OD_600_ of 0.5 and then diluted 20-fold into new tubes containing fresh identical medium containing the same concentration of glucose and acetate. Growth rates were determined by measuring optical density between OD_600_ 0.05 and 0.5 after dilution. Remarkably, we observed that cultures grown initially on glucose exhibited about 25% higher growth rates on the mixed substrates (Figure 1B and Figure S1). Performing several dozen repeats of this experiment, we did observe this phenomenon to be somewhat fickle and in around 30% of biological repeats the state was disrupted, and the two cultures very quickly converged to the same growth rates. However, in the remainder of experiments shown in Fig. 1B, we observed a significant difference in growth rates that persisted as long as we followed the cultures, for at least 6 generations. This growth rate difference was much larger than what would be expected naively from existing models^4^ and based on simple dilution of the existing proteome. Because growth rate on acetate alone is 50% slower than growth on glucose alone, at the beginning of the growth curve, at OD_600_ 0.05, one would naively expect a 10% difference in growth rates, while at the end of the growth curve, at OD_600_ 0.5, the difference should be 1%. To determine if these divergent growth rates were perhaps converging slowly, we diluted the cultures three consecutive times in identical growth medium and measured growth rates again. Remarkably, the distinct, history-dependent growth rates persisted for at least 15 generations, although we cannot rule out very slow convergence with additional rounds of dilution (Fig. 1C). Conservatively, estimating the dilution of initial proteome composition, at the beginning of the growth curve of the third dilution, the proteome from pre-shift conditions should be diluted 2000-fold, hence there is no reason to expect any detectable differences in growth rates at this point.

**Figure 1:**
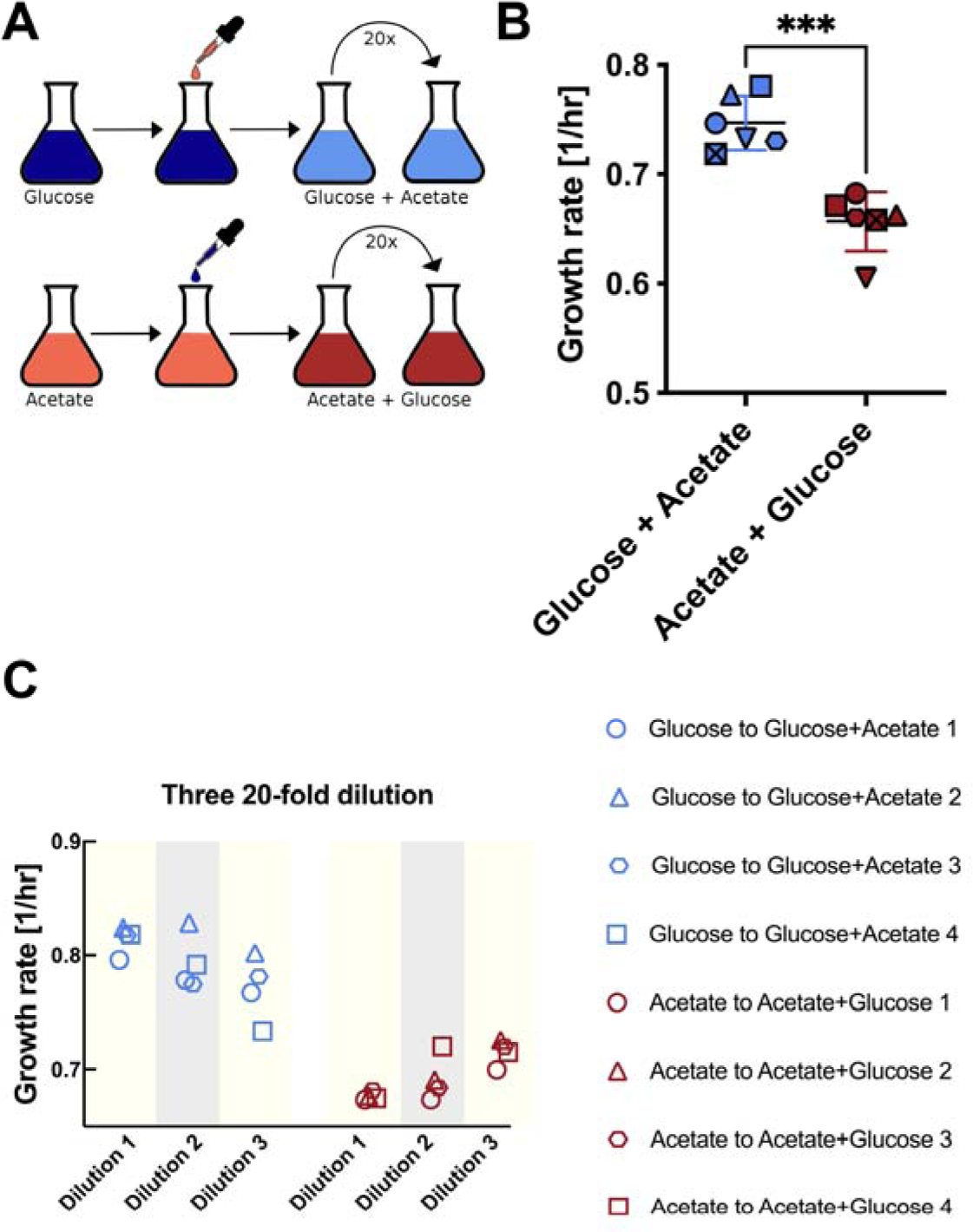
Shifts from single to two carbons sources leads to history-dependent growth rates. **A**, Experimental procedure: All growth measurements were done at 37°C in 500 ml flasks shaking at 220 rpm. *E. coli* NCM3722 was inoculated from a single colony from LB plate and grown in LB medium. The culture was then split to N^+^C^+^ minimal medium with either glucose or acetate as the single carbon source and grown overnight. On the next day, the cultures were transferred to fresh N^+^C^+^ minimal medium with either 20 mM glucose or 60 mM acetate and first grown for 5-6 generations. Then in mid-exponential phase (roughly at OD_600_ 0.2), glucose was added to the acetate culture and acetate was added to the glucose culture, to make an identical final concentration of 20mM glucose and 60mM acetate in both tubes (carbon concentrations were high, such that carbon consumed by growth could be neglected). When the cultures reached an OD_600_ of 0.5, they were diluted 20-fold. The growth rates of the diluted cultures were determined by taking at least four OD_600_ measurements between OD_600_ 0.05 and OD_600_ 0.5. **B**, Growth rates of *E. coli* with different growth history growing in the same medium, containing both glucose and acetate, sampled and calculated in the 20-fold diluted cultures as described in A. The symbols indicate biological repeats that were run in parallel and can be compared for different growth histories. The colors indicate the direction of the upshift (red = start from acetate only, blue = start from glucose only). Error bars are standard deviations of growth rates. **C**. We monitored the growth of the culture for three consecutive rounds of 20-fold dilutions. We found that the differences between the growth states persisted, though we cannot rule out very slow convergence of the growth rates.

Next, we investigated whether this history-dependence was specific to glucose and acetate or a more general phenomenon. To address this question, we first tested several combinations of glucose with other gluconeogenic organic acids (Fig. S1). Indeed, we found history-dependence for combinations of glucose with gluconeogenic substrates other than acetate, including lactate, pyruvate and succinate. We also observed that growth rates on many combinations of glucose with gluconeogenic carbon sources were substantially slower than growth rates on glucose alone (Fig. S2). Interestingly, when testing different carbohydrates in combination with acetate (Fig. S1), we observed history-dependent growth rates for combinations of phosphotransferase system (PTS) transported carbohydrates with acetate, while non-PTS sugar lactose had no history dependence. Strikingly, for the fructose-acetate and mannose-acetate combinations, the order of the growth rates was reversed, with the culture coming from acetate growing faster. Our results indicate that history-dependent growth states are not specific to the glucose-acetate combination, but instead occur for diverse combinations of substrates.

To determine if the difference in growth rates between the two hysteresis states was also reflected in metabolism, we characterized hysteresis states for different carbon combinations by measuring absolute concentrations of 25 central metabolites by LC-MS/MS^13^ and, to obtain flux information, measuring dynamic ^13^C-labeling patterns of intracellular metabolite pools (Figs. S3-S9) and labeling patterns of proteinogenic amino acids by GC-MS (Fig. S10). In these experiments, addition of either labeled glucose or acetate to both hysteresis cultures, resulted in a new steady-state within 20s indicating rapid utilization of both substrates. Consistent with the different growth rates, we found levels of about half the detectable metabolites in central metabolism to differ by up to a factor two. Surprisingly, many TCA metabolites (citrate, fumarate, malate) were elevated in the faster growing state that was first grown on glucose (Fig. S4).

Since history-dependence requires different proteomes, we tested whether we could disrupt it by knockout of transcriptional regulators and enzymes, which could be involved in mediating the positive feedback required for bistability or by changing initial conditions. Guided by the pattern of carbon sources that showed history-dependence, we focused on transcription factors related to the phosphotransferase system, gluconeogenesis, and acetate metabolism (see Fig. 2A & Fig. S11). Indeed, we found that the phenomenon was abolished in a knockout of the transcriptional regulator Mlc (Fig. 2B), which controls the expression of enzymes of the PTS systems^14,15^. Consistently, the Mlc knockout also abolished hysteresis for the combinations of mannose and acetate (Fig. 2B). Theoretical studies previously proposed that the autocatalytic nature of the PTS system could give rise to bistability^16^.

**Figure 2:**
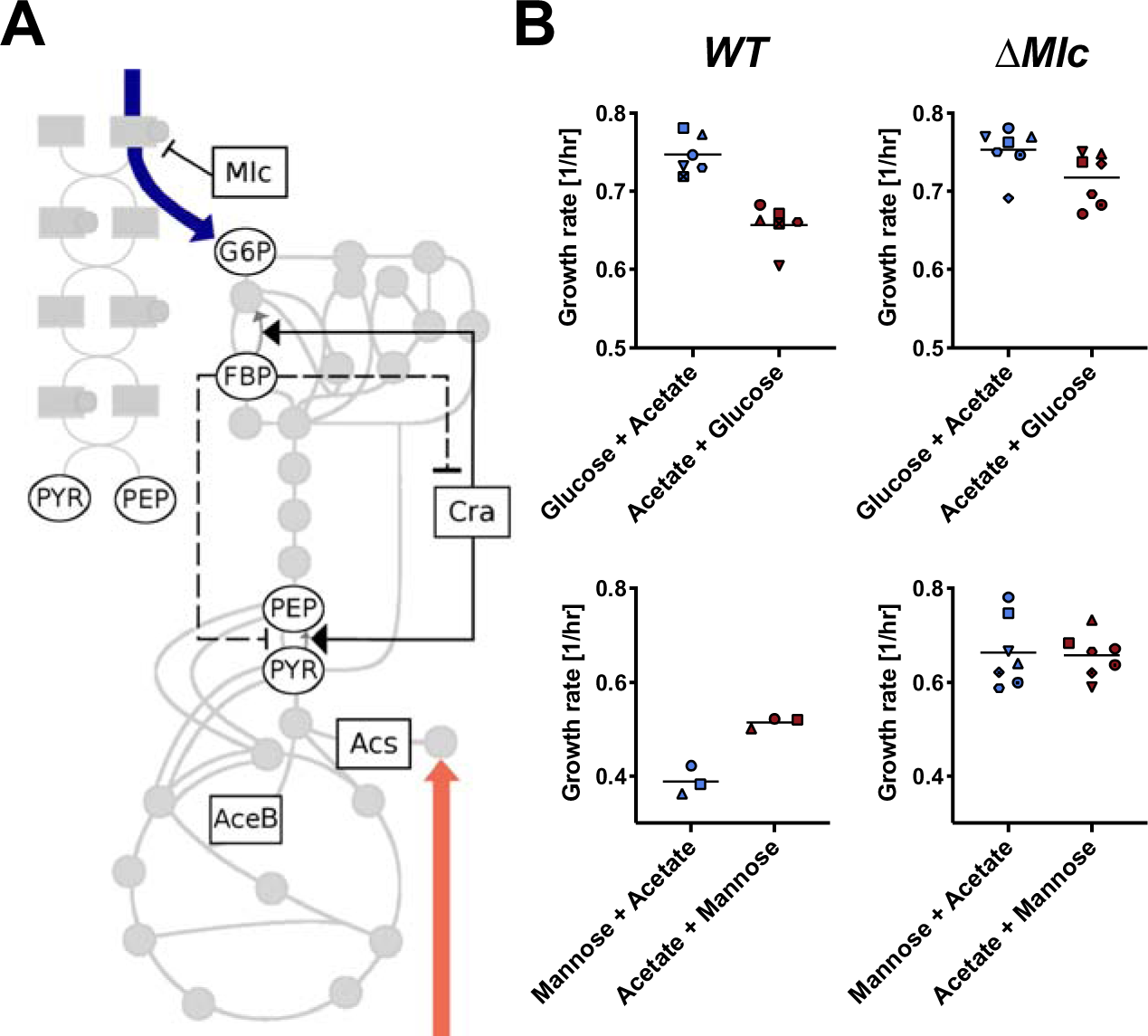
Knockout of the transcription factor Mlc abolished growth rate hysteresis. **A**, Schematic illustration of central carbon metabolism, the phosphotransferase system (PTS). Key transcriptional regulators and enzymes involved in glucose and acetate metabolism, as well as gluconeogenesis and PTS are highlighted. We measured growth rate hysteresis in knockout strains of these genes, as well as using inducible genetic constructs when these genes were essential for growth on acetate (Fig. S11). **B**, Growth rate hysteresis was abolished in an Mlc knockout strain for glucose and acetate. We also tested the mannose and acetate carbon combination and found that indeed, hysteresis was also abolished in an Mlc knockout. Mlc is the major transcriptional regulator of the PTS system. For comparison, WT glucose/acetate is shown (same as Fig. 1B). The symbols indicate biological repeats that were run in parallel and can be compared for different growth histories.

While it was previously known that history-dependent states in microbes can persist for a few generations during medium shifts^4^ and affect developmental processes like sporulation^6^, here we find that central carbon metabolism can directly result in long-term cellular memory. A striking feature of growth on some combinations of substrates (e.g. glucose&acetate, glucose&lactate, glucose&pyruvate, lactose&acetate) is that growth rate is slower than on the faster substrate alone. It is possible that this could provide an advantage for bacteria in fluctuating environments^6,17,18^: for example when acetate is present for long periods and glucose only during short pulses. Not fully adapting cellular metabolism to glucose would in this case be advantageous because it would reduce the time needed to switch back to growing on acetate alone.

To test this hypothesis, we shifted the cultures with two different histories to medium containing acetate as the sole carbon source. We quantified the rate of adaptation in each case by measuring lag times. The difference in lag times between the two cultures was small and difficult to resolve. However, both cultures showed a reduced lag time on acetate as compared to bacteria growing at a faster rate on glucose only^19^ (Fig. 3). Hence, while slower growth rates on combinations of substrates, as compared to single substrates may seem counterintuitive, one benefit is that they can enable faster adaptation to the non-preferred substrate if the preferred substrate runs out. Therefore, reduced growth rates on combinations of glycolytic and gluconeogenic substrates could be a bet-hedging strategy, rather than a regulatory malfunction.

**Figure 3:**
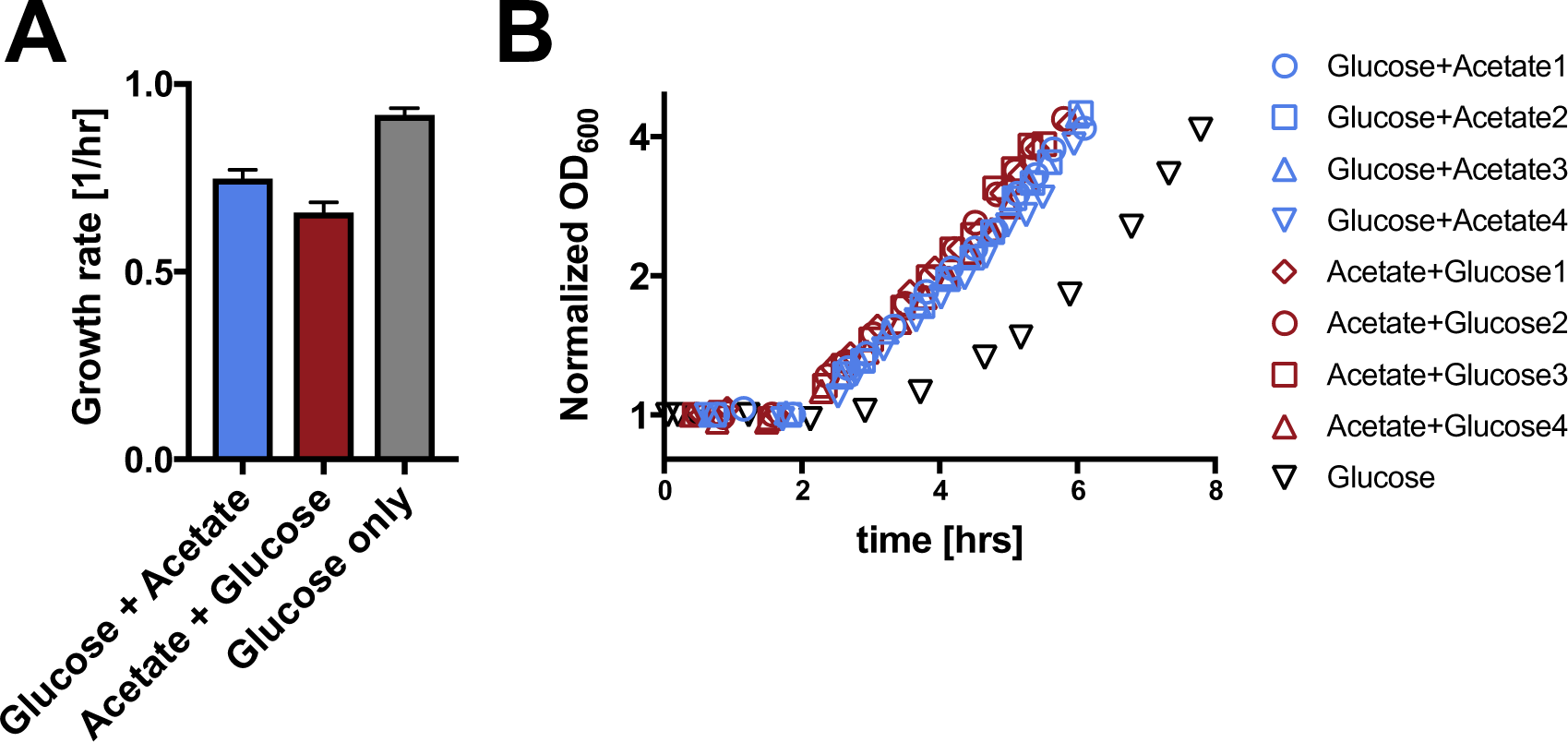
Reduced growth rates on carbon combinations result in faster switching to the non-preferred substrate. **A**, Growth rates of the two hysteresis states growing on the combination of glucose and acetate, versus growth rates on glucose alone. (Same data as Fig. 1B, 2 biological replicates for glucose only). All error bars are standard deviations. **B**, Bacteria from the three growth states in A were filtered and resuspended in minimal medium containing acetate as the sole carbon source. Growth resumption was measured as a function of time after the shift to acetate and OD_600_ was normalized to the initial value. The figure shows four independent biological repeats. The time course for the glucose only condition was taken from a parallel work^19^. Strikingly, while growth rates on the combination of glucose and acetate were slower compared to glucose only medium, these slower growth rates enabled substantially faster switching to the non-preferred substrate acetate.

## Acknowledgements

We thank Maren Diether and Elad Noor for fruitful discussions and helpful suggestions. This project was supported by MIRA grant (5R35GM137895) and an HMS Junior Faculty Armenise grant to MB and an IPhD fellowship of SystemsX.ch to D.C.

