## supplementary information for "Long-term history dependence of growth rates of *E. coli* after nutrient shifts"

Christodoulou et al.

### Supplementary Information

### Materials and Methods

#### Bacterial cultures

All experiments were performed with *Escherichia coli* NCM3722 (from now on referred to simply as *E. coli*). The gene knockouts used in the genetic perturbation experiments were derived from the KEIO collection [1] and introduced into this strain via P1-phage transduction. Strains with genes inducible by chlortetracycline (cTc) (pTet-*tetR*/pTet-*Cra*, pTet-*tetR*/pTet-*AceB*) were constructed as described before [2]. Strains under control of 3-Methylbenzyl were previously described in You et al. [3] and were kindly shared by the lab of Terry Hwa. Frozen glycerol stocks were grown on lysogeny broth (LB) agar plates. From there, 3 ml of liquid LB medium were inoculated with a single colony and after 5-8 hours incubation on a shaker at 37°C and 250 rpm, 3 ml of $N^-C^-$ minimal medium (23 mM $K_2SO_4$, 310 mM $K_2HPO_4$, 140 mM $KH_2PO_4$, 15 mM $MgSO_4$, 170 mM NaCl, 20 mM $NH_4Cl$) [4,5] supplemented with one or two carbon sources (sugars or organic acids, each at a concentration corresponding to 120 mM atomic carbon), were inoculated with the LB preculture at 500 to 1000 fold dilution and incubated at 37°C and 250 rpm overnight. For the main experiment, cultures were grown in 500 mL shake flasks containing 40 mL of the medium described above and were incubated on a shaker (250 rpm) at 37°C.

#### Carbon shift experiment

Minimal medium ($N^-C^-$) [4,5] containing either one glycolytic or one gluconeogenic carbon was inoculated with an overnight culture of *E. coli* pre-grown on the same medium. Cultures were grown to an optical density ($OD_{600}$) of 0.2. Then, to the culture growing on only glycolytic carbon source, the gluconeogenic carbon source was added and vice versa so that the final concentration of both carbon sources was the same in either case (that is, they have the same final concentration of atomic carbon as explained in the Bacterial cultures section, either when starting with the glycolytic carbon first or with the gluconeogenic). Upon reaching $OD_{600}$ of 0.5, the cultures were diluted 20-fold into medium containing again both carbon sources. The growth rate was afterwards determined by manually taking at least four readings of $OD_{600}$. By fitting a line to the logarithm of these $OD_{600}$ measurements versus time, we determined growth rates. The experimental procedure is outlined in Figure 1 A. When performing this experiment with strains expressing genes essential for growth on acetate under the control of the Tet promoter, cTc was added to medium containing acetate as sole carbon source at a concentration necessary to achieve wild type growth rate. After addition of the second carbon, the inducer was removed by a washing step. Cultures were then diluted into two different flasks containing $N^-C^-$ supplemented with both carbon sources. To one of these, cTc was added at the same concentration as previously described.

#### Metabolomics and $^{13}$C-labeling

Metabolite samples were taken at the end of the experiment described in the previous section, when the cultures reached $OD_{600}$ of 0.5 after the dilution step. Sampling, sample processing and liquid-chromatography mass spectrometry (LC-MS/MS) measurements were performed according to a previously published method [6]. Dynamic labeling samples were taken at the same time, following the protocol published by [7]. Two differently labeled media were used: (A) $N^-C^-$ supplemented with 100% U-$^{13}$C-acetate plus 100% unlabeled glucose, (B) $N^-C^-$ supplemented with 100% unlabeled acetate, 50% unlabeled glucose and 50% U-$^{13}$C-glucose (see Figure 2A). Samples were analyzed by LC-MS. In order to obtain steady state labeling data, when performing the dilution step of the carbon shift experiment described in the previous section, cultures were diluted into labeled medium A and B as described above. In order to remove any unlabeled medium, cells were first transferred to a cellulose filter on a Buchner flask attached to a vacuum pump. Subsequently, the filter was transferred to a flask containing either labeled medium A or labeled medium B. Cultures were grown to an $OD_{600}$ of 0.5, then metabolite samples were taken as described above. In addition, cell pellets were harvested for GC-MS analysis as described in [8] and [9].

**Processing and analysis of MS data**

From LC-MS data, metabolite abundances were calculated using the in-house tool mssoftware. The obtained isotopomer abundances from the dynamic labeling experiment were analyzed using MATLAB and R, following a similar procedure as presented in Yuan *et. al*, 2009 [10]. Isotope fractions from GC-MS data were obtained using FiatFlux [11].

#### Supplementary Figures

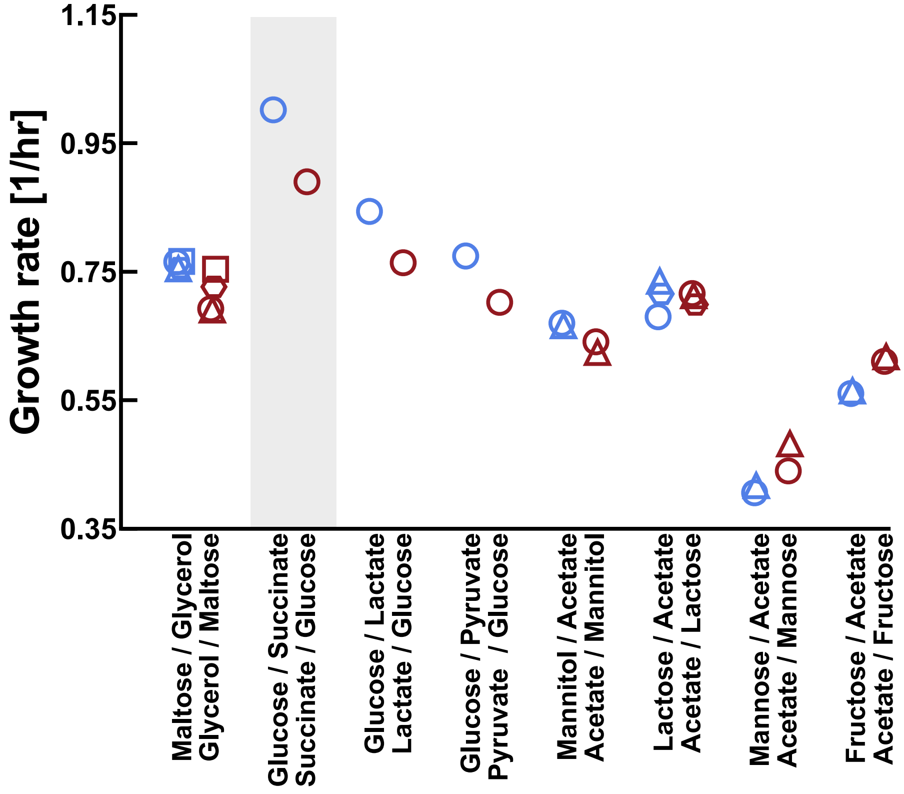

.

**Figure S1: Hysteresis experiment with other carbon combinations.** The same experiment described in Fig. 1A, was repeated for other combinations of carbon source. Different symbols indicate specific biological repeats. We indeed found history dependent growth rates for several other substrate combinations.

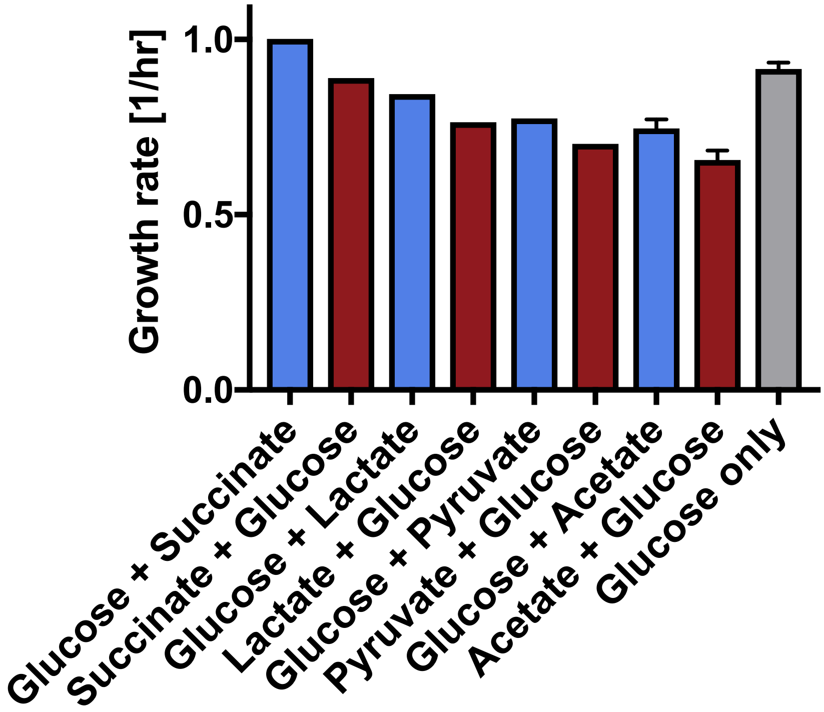

**Figure S2: Growth rates of hysteresis states for carbon combinations with glucose versus growth rate on glucose alone.** Growth rates on many carbon combinations were slower than on glucose alone. Glucose and succinate were an exception.

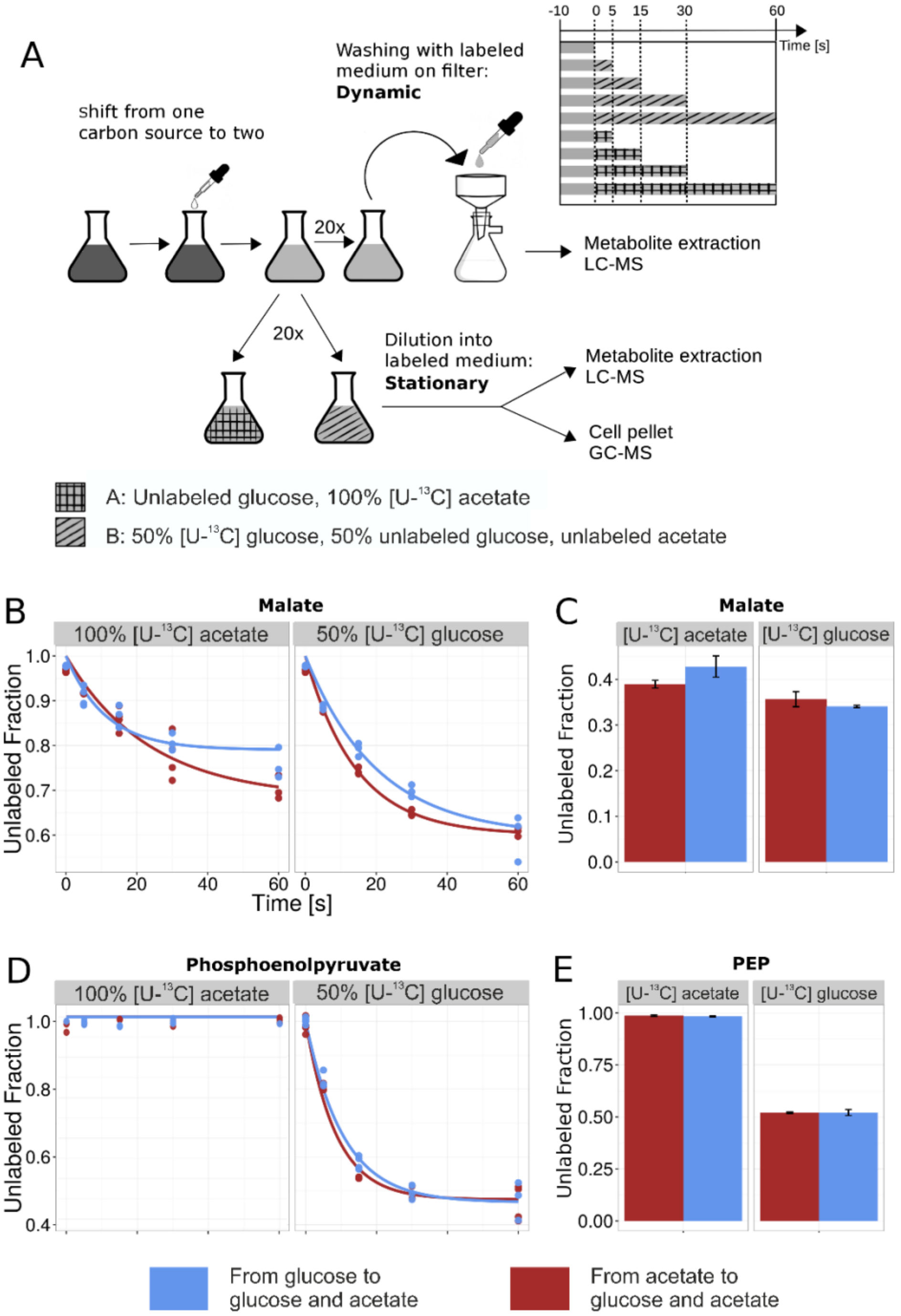

**Figure S3: Illustrative examples from metabolomics and $^{13}$C-labeling experiments to elucidate metabolic state of *E. coli* during hysteresis. A,** Schematic representation of the experimental procedure. *E. coli* was first shifted from one carbon source (either glucose or acetate) to both. In mid-exponential phase, cultures were diluted into unlabeled medium (top, dynamic labeling) or medium supplemented with either [U-$^{13}$C] acetate or [U-$^{13}$C] glucose (bottom, different labels are shown as checked (A, acetate) or striped (B, glucose)). Samples for stationary labeling were taken after growing the cultures for several hours on labeled medium. In order to estimate labeling dynamics, cells were washed with labeled medium A or

B on a cellulose filter for different durations as illustrated in the scheme at the top right. **B,** Labeling dynamics in malate. Shown is the decay of the unlabeled fraction over time in cases where cells were washed with medium containing [U-$^{13}$C] acetate (left panel) or [U-$^{13}$C] glucose (right panel). Points are individual measurements. The lines are obtained from a non-linear least squares fit of an exponential decay function to the data. The colors correspond to the direction of the carbon shift. **C**: Stationary labeling pattern in malate. Shown is only the unlabeled fraction. **D**: Labeling dynamics of phosphoenolpyruvate (PEP). **E**: Stationary labeling pattern in PEP.

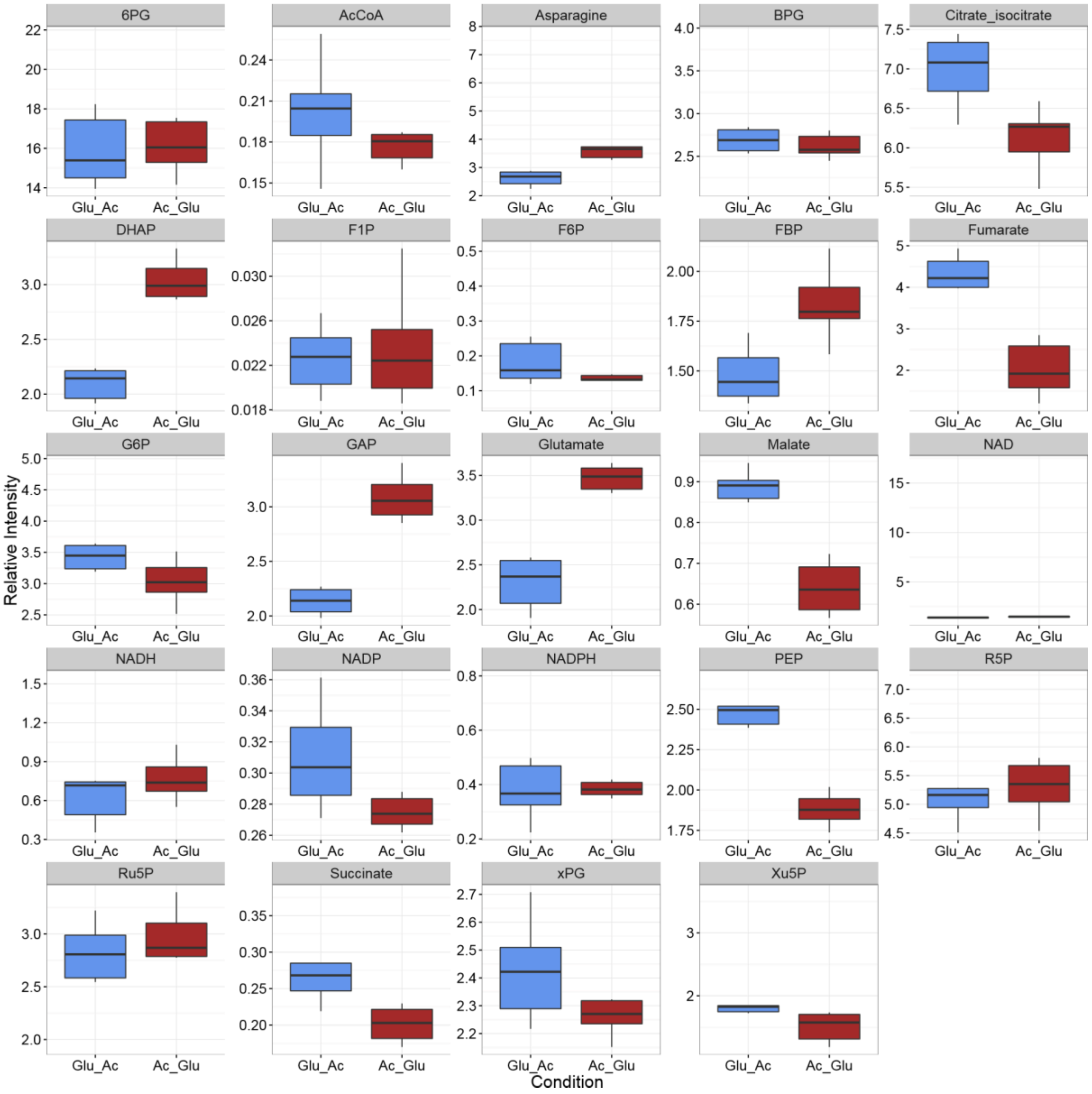

**Figure S4: Comparison of metabolite pools between conditions.** Shown are for each measured metabolite the relative intensities (12C signal / 13C internal standard) for shifts from glucose (blue) or acetate (red) to glucose and acetate.

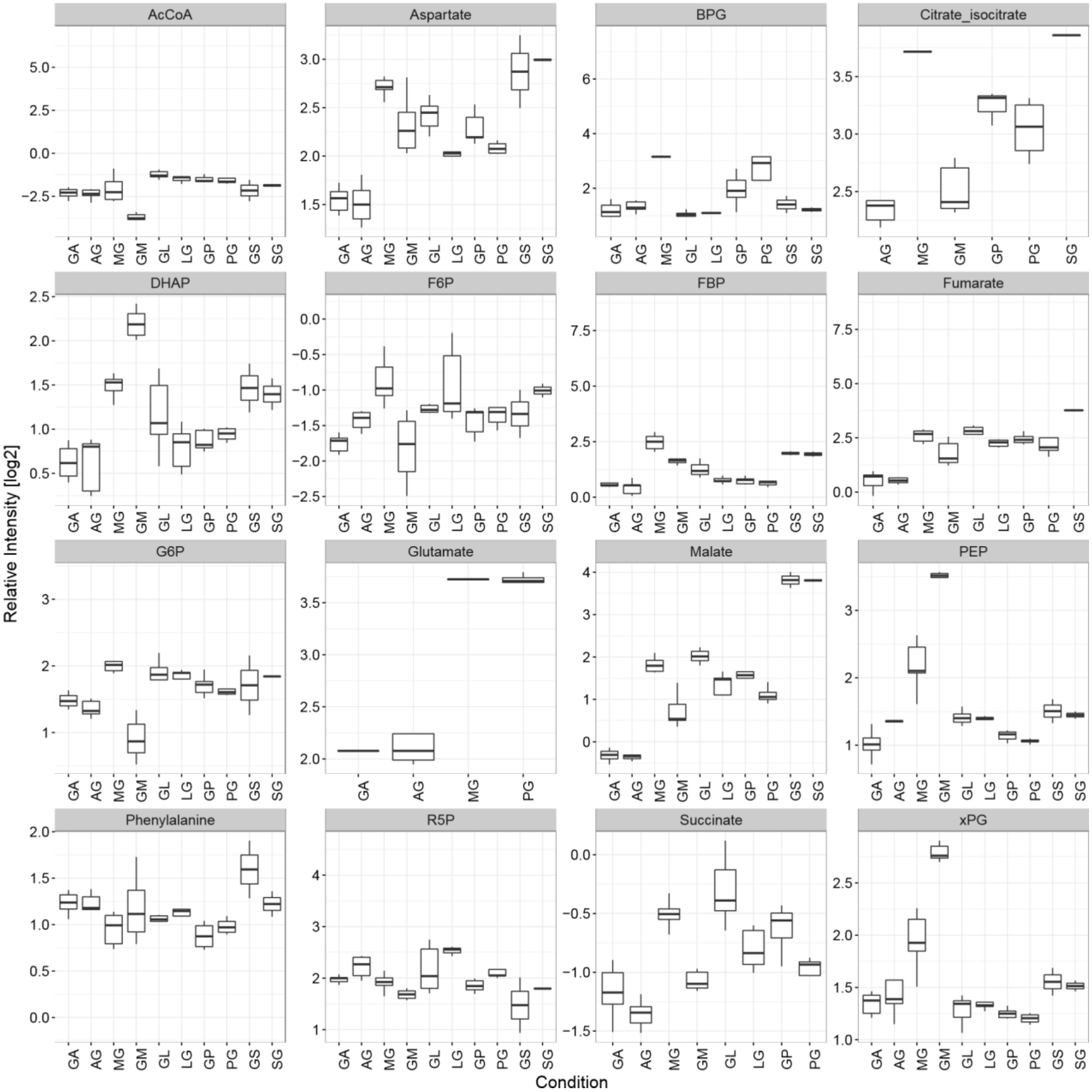

**Figure S5: Comparison of metabolite pools between conditions**. Shown are for each measured metabolite the relative intensities (12C signal / 13C internal standard) for different carbon shifts (GA = glucose + acetate, AG = acetate + glucose, MG = maltose+ glycerol, GM = glycerol + maltose, GL = glucose + lactate, LG = lactate + glucose, GP = glucose + pyruvate, PG = pyruvate + glucose, GS = glucose + succinate, SG = succinate + glucose).

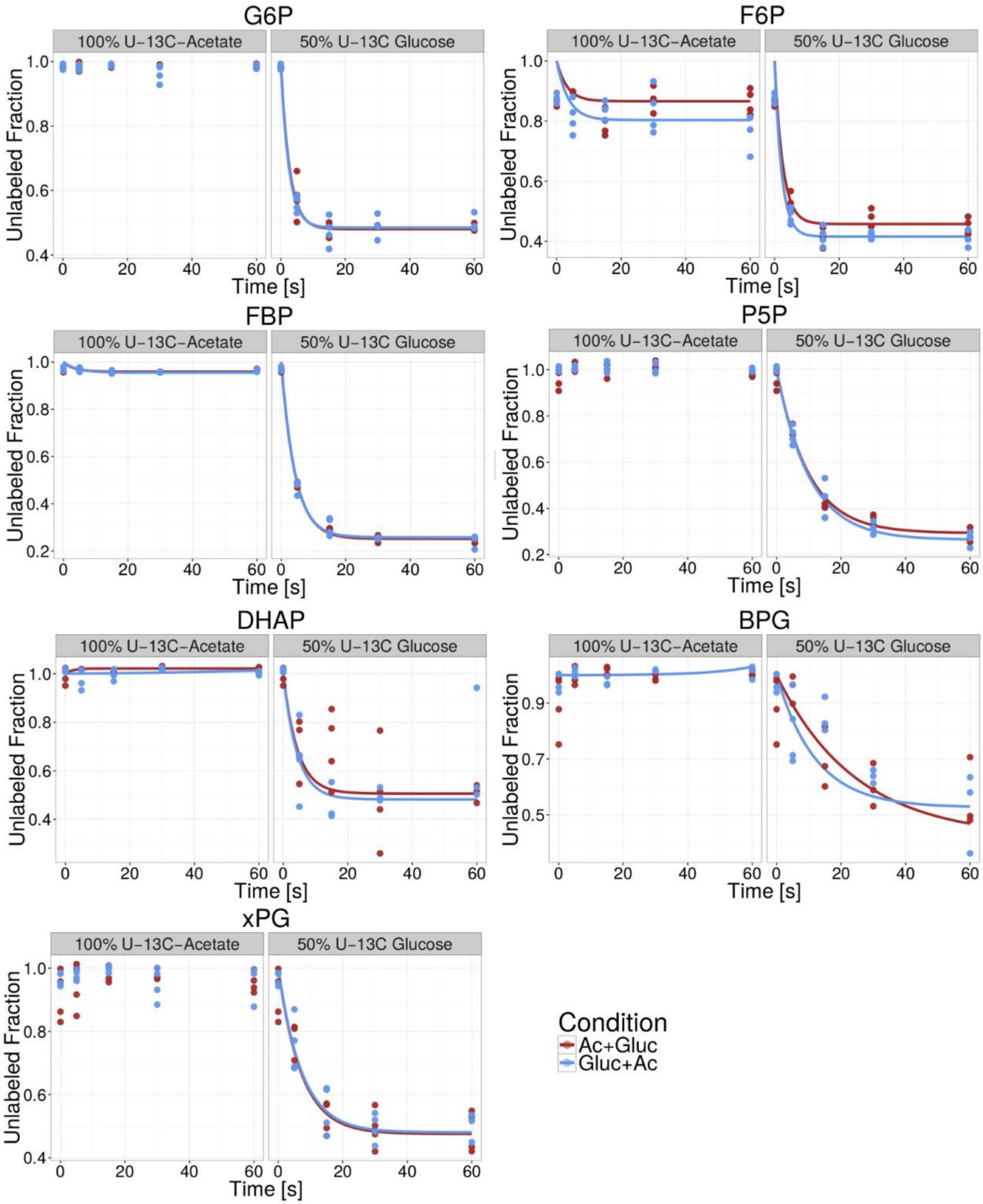

**Figure S6: Labeling dynamics for different metabolites in glycolysis**. Shown is the decay of the unlabeled fraction over time in cases where cells were washed with medium containing labeled acetate (left panel) or labeled glucose ( right panel). Points are individual measurements. The lines are obtained from a non-linear least squares fit of an exponential decay function to the data.

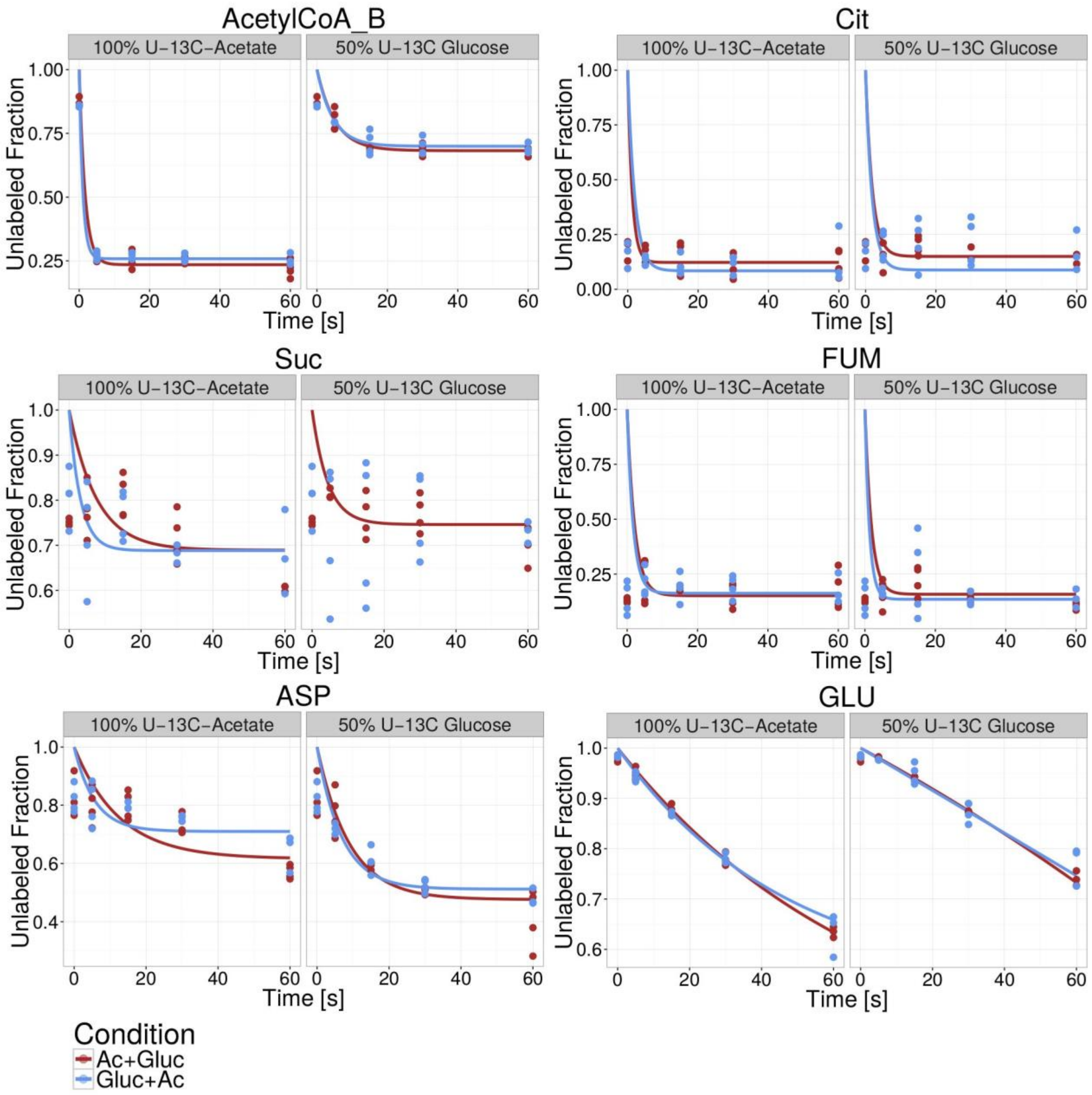

**Figure S7: Labeling dynamics for different metabolites of the TCA cycle**. Shown is the decay of the unlabeled fraction over time in cases where cells were washed with medium containing labeled acetate (left panel) or labeled glucose (right panel). Points are individual measurements. The lines are obtained from a non-linear least-squares fit of an exponential decay function to the data. The colors correspond to the direction of the carbon shift. The very low unlabeled fraction in citrate and fumarate at time zero can be explained by overall very low intensity and noisy signal.

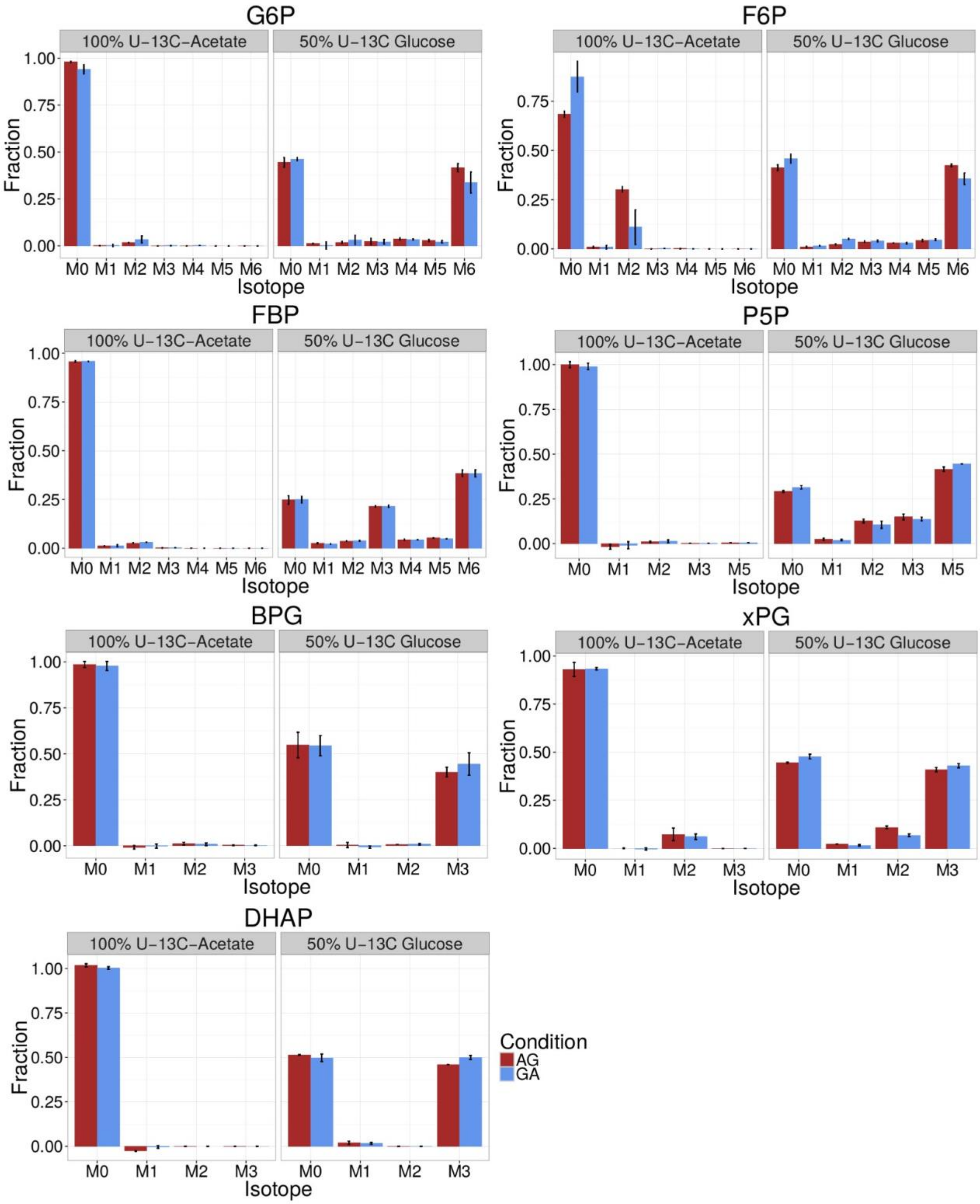
G6P
F6P
FBP
P5P
BPG
xPG
DHAP
100% U-13C-Acetate
50% U-13C Glucose
Fraction
Isotope
M0
M1
M2
M3
M4
M5
M6
0.00
0.25
0.50
0.75
1.00
Condition
AG
GA

**Figure S8: Steady state labeling pattern of different metabolites in glycolysis as determined by LC-MS.** Bars indicate mean fractions of different mass isotopomers. Error bars are standard deviations (n=4). The colors correspond to the direction of the carbon shift.

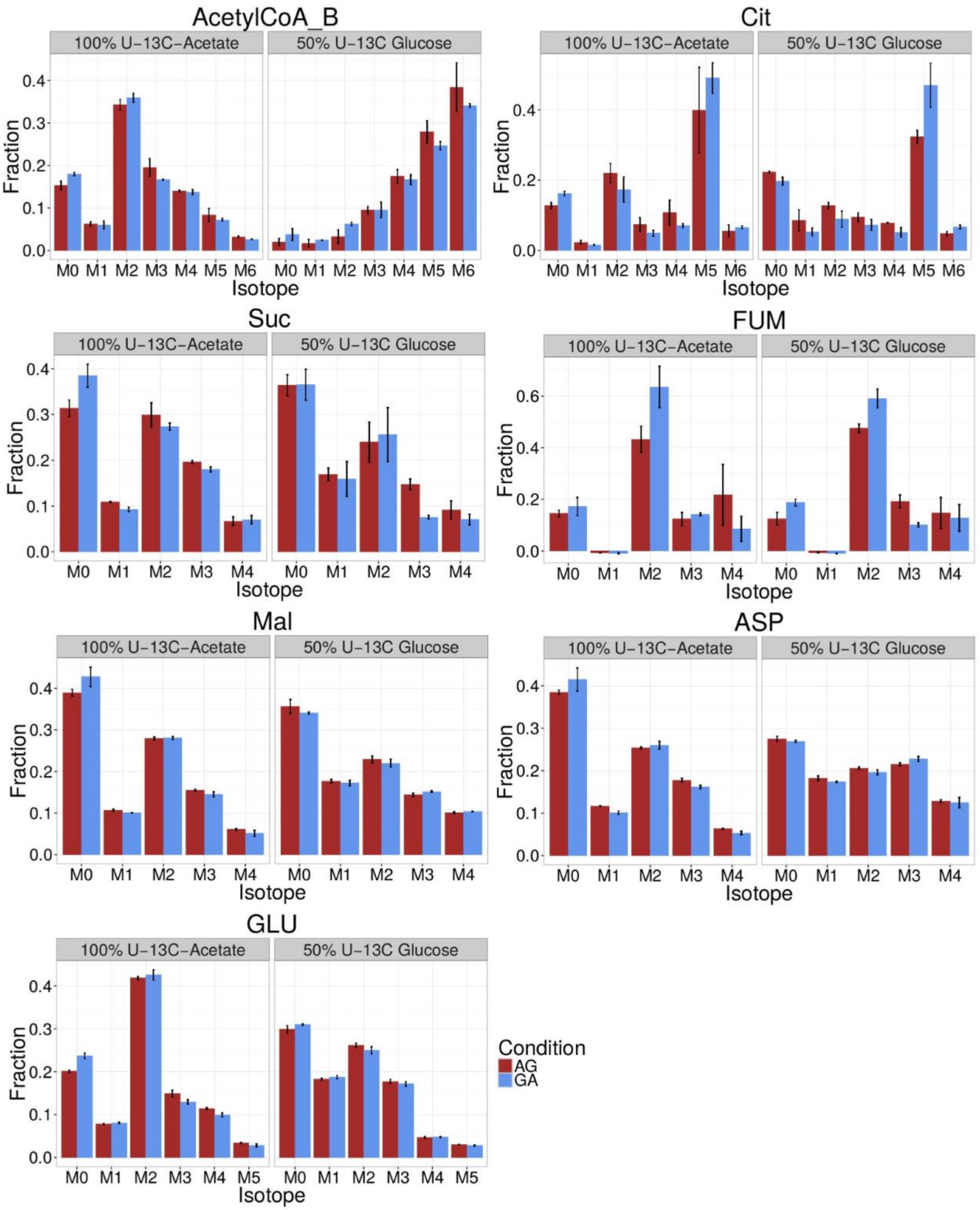
AcetylCoA_B
Cit
Suc
FUM
Mal
ASP
GLU
100% U-13C-Acetate
50% U-13C Glucose
Fraction
Isotope
M0 M1 M2 M3 M4 M5 M6
Condition
AG
GA

**Figure S9: Steady state labeling pattern of different metabolites in the TCA cycle as determined by LC-MS**. Bars indicate mean fractions of different mass isotopomers. Error bars are standard deviations (n=4). The colors correspond to the direction of the carbon shift.

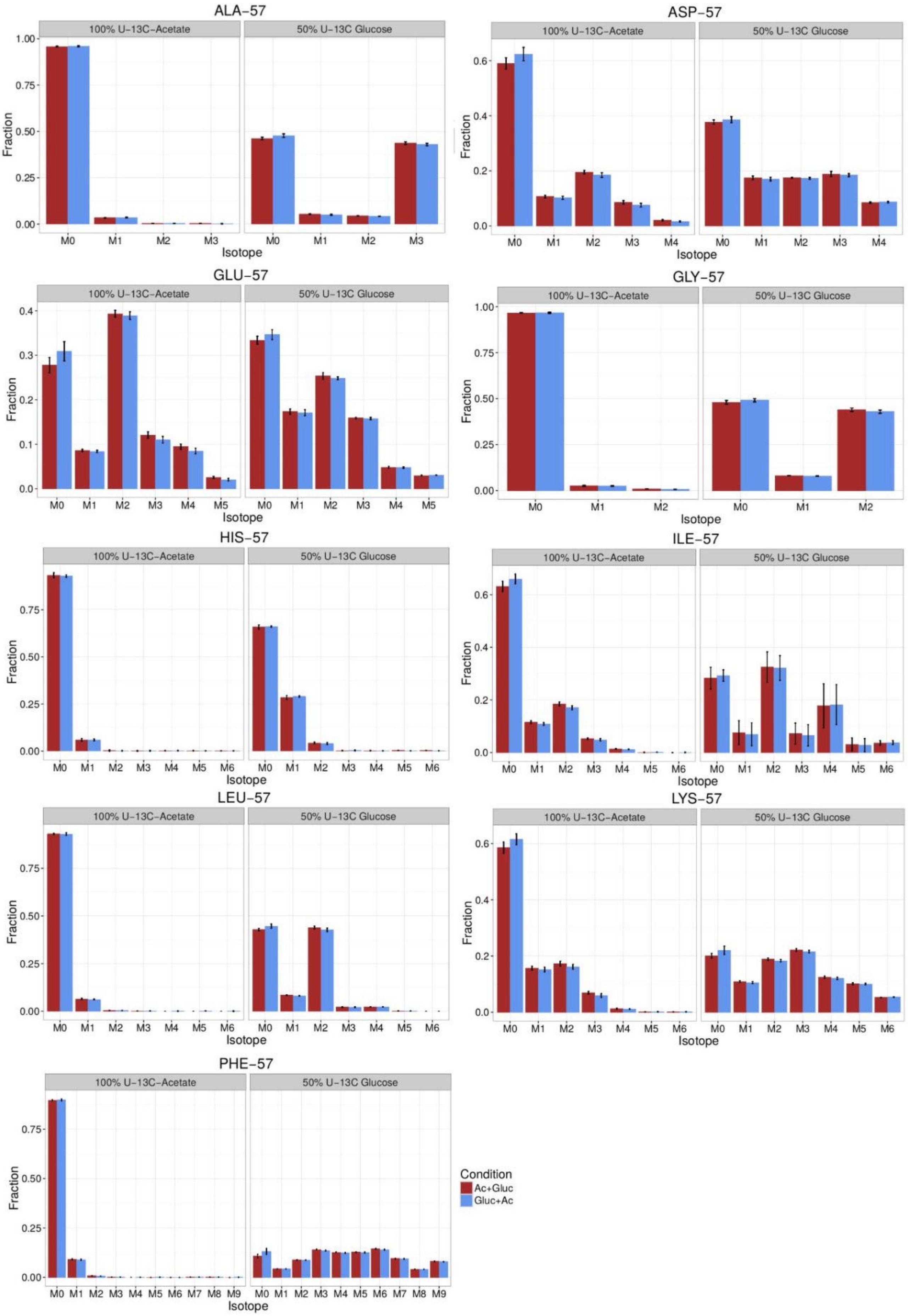
ALA-57
ASP-57
GLU-57
GLY-57
HIS-57
ILE-57
LEU-57
LYS-57
PHE-57
100% U-13C-Acetate
50% U-13C Glucose
Fraction
Isotope
M0
M1
M2
M3
M4
M5
M6
M7
M8
M9
Condition
Ac+Gluc
Gluc+Ac

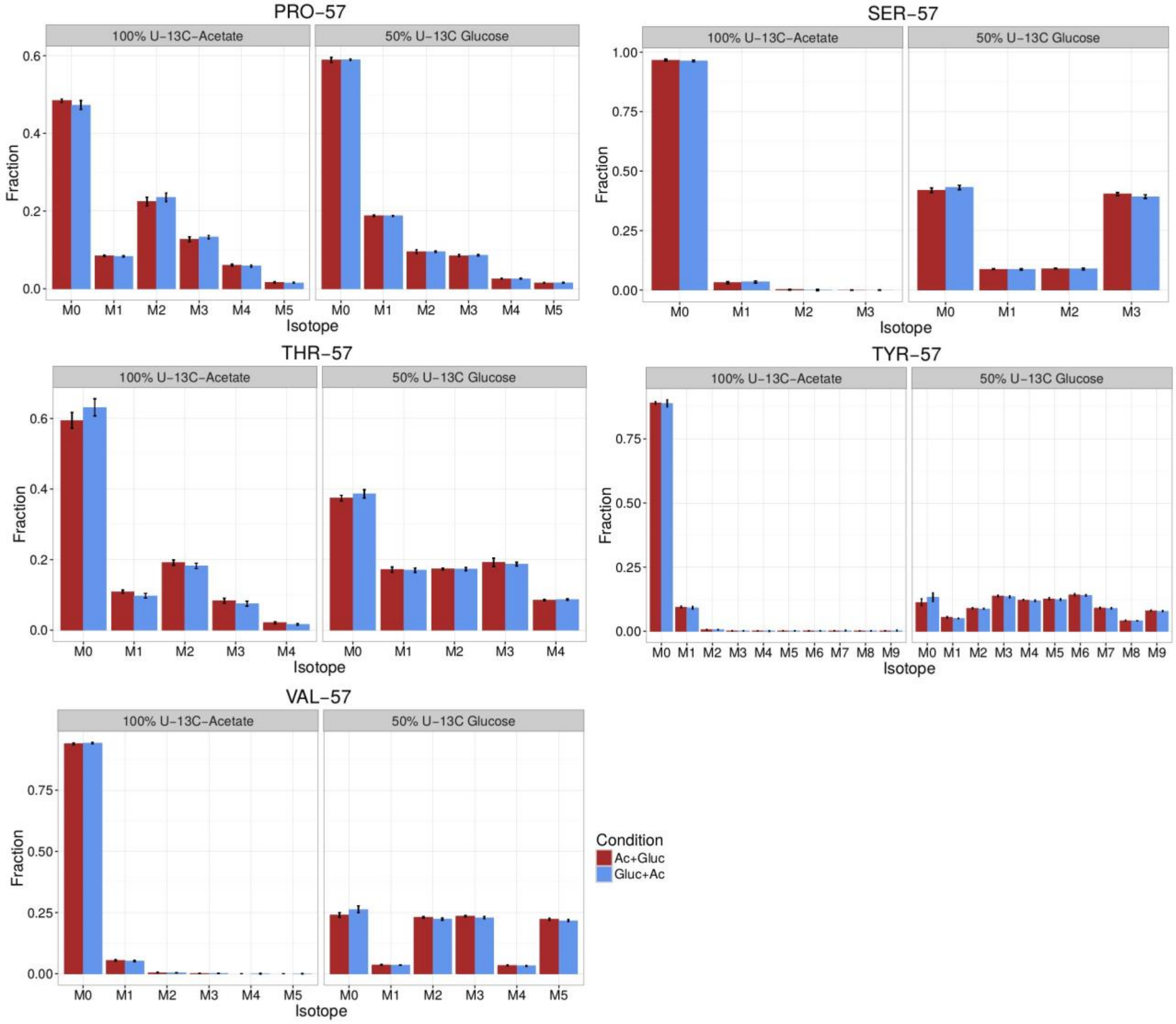

**Figure S10: Steady-state labeling pattern of proteinogenic amino acids as determined by GC-MS**. The data obtained show the precursor ion (amino acid derivatized with tert-Butyldimethylsilane) minus a tert-butyl group (m/z=57). Bars indicate mean fractions of different mass isotopomers. Error bars are standard deviations (n=4). The colors correspond to the direction of the carbon shift.

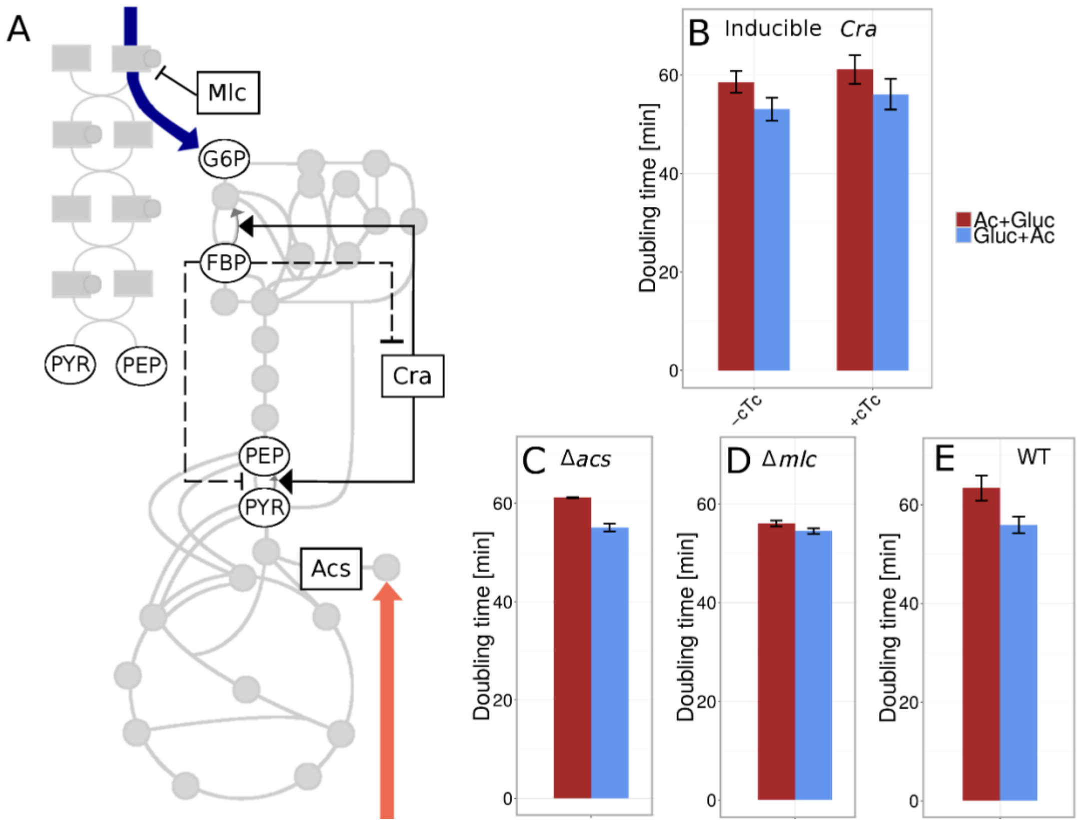

**Figure S11: Doubling times on glucose + acetate and acetate + glucose for *E. coli* with different genetic background**. **A**: Schematic overview of the central carbon metabolism with proteins that could potentially influence the hysteresis, highlighted. Boxes represent proteins, circles metabolites, and boxes with attached circles phosphorylated proteins. The blue and orange arrows correspond to the influx of glucose and acetate respectively. Regulatory interactions are shown by solid (transcriptional) and dashed (allosteric) lines. **B**: Doubling times of an *E. coli* strain expressing the transcriptional regulator Cra under the control of an inducible promoter. Significantly different doubling times depending on the direction (two-sample t-Test p-value = 0.026 in the absence of inducer, and p-value = 0.02 in the presence of inducer) **C**: Doubling times of an *E. coli* Δacs strain. Significantly different doubling times depending on the direction (two-sample t-Test p-value = 0.008) **D**: Doubling times of an *E. coli* Δmlc strain. Non-significant difference in doubling times depending on the direction (two-sample t-Test p-value = 0.461) **E**: Doubling times of an *E. coli* WT strain. Significantly different doubling times depending on the direction (two-sample t-Test p-value = 0.0185). All plots show weighted average doubling times. Colors indicate the direction of the carbon upshift. Error bars represent standard deviations. The numbers of replicates are B: 9, C: 6 , D: 5, E: 6.
